# Sound localization of world and head-centered space in ferrets

**DOI:** 10.1101/2021.09.15.460425

**Authors:** Stephen M. Town, Jennifer K. Bizley

## Abstract

The location of sounds can be described in multiple coordinate systems that are defined relative to ourselves, or the world around us. Evidence from neural recordings in animals point towards the existence of both head-centered and world-centered representations of sound location in the brain; however, it is unclear whether such neural representations have perceptual correlates in the sound localization abilities of non-human listeners. Here, we establish novel behavioral tests to determine the coordinate systems in which ferrets can localize sounds. We found that ferrets could learn to discriminate between sound locations that were fixed in either world-centered or head-centered space, across wide variations in sound location in the alternative coordinate system. Using probe sounds to assess broader generalization of spatial hearing, we demonstrated that in both head and world-centered tasks, animals used continuous maps of auditory space to guide behavior. Single trial responses of individual animals were sufficiently informative that we could then model sound localization using speaker position in specific coordinate systems and accurately predict ferrets’ actions in held-out data. Our results indicate that an animal model in which neurons are known to be tuned to sound location in egocentric and allocentric reference frames can also localize sounds in multiple head and world-centered spaces.

**Significance Statement:** Humans can describe the location of sounds either relative to themselves, or in the world, independent of their momentary position. These different spaces are also represented in the activity of neurons in animals, but it’s not clear whether non-human listeners also perceive both head and world-centered sound location. Here, we designed two behavioral tasks in which ferrets had to discriminate between two sounds using their position in the world, or relative to the head. Subjects learnt to solve both problems and showed the ability to generalize sound location in each space when presented with infrequent probe sounds. These findings reveal a perceptual correlate of neural sensitivity previously observed in the ferret brain and establish that, like humans, ferrets can access an auditory map of their local environment.

## Introduction

The ability to localize sounds in our environment is critical for both humans and other animals, across a variety of behaviors and in a wide range of ecological settings. In contrast to other senses such as vision or somatosensation, the auditory system lacks a spatially-organized sensory epithelium such as the retina or skin from which to extract spatial information, and so sound location must be computed within the brain. Extensive research has shown how sound localization cues are extracted in the midbrain (Tollin and Yin, 2002; Yin and Chan, 1990) and transmitted across the brain to areas, including auditory cortex (Keating et al., 2015; Stecker et al., 2005), parietal and prefrontal cortex (van der Heijden et al., 2019). While midbrain neurons show tuning to specific localization cues, auditory cortex contains cue-invariant representations of sound location (Wood et al., 2019) and is essential for sound localization in primates and carnivores (Slonina et al., 2022).

The study of spatial coding in the auditory system has often tacitly assumed that sound location is encoded in head-centered space. However, emerging evidence suggests that sound location may be represented in multiple coordinate frames across the auditory system. These include egocentric coordinate systems centered on body parts such as the eyes (Andersen and Buneo, 2002; Caruso et al., 2021; Groh et al., 2001; Werner-Reiss et al., 2003) and world-centered (or *allocentric)* coordinates that map sounds into a listeners’ environment across changes in head position and direction (Amaro et al., 2021; Town et al., 2017). Generating world-centered representations of sound location requires that neural circuits involved in the spatial analysis of sound compensate for effects of head rotation by listeners. Indeed, cells in the dorsal cochlear nucleus integrate auditory and vestibular information, offering a potential mechanism to discriminate self and source motion (Wigderson et al., 2016; Wu and Shore, 2018). Yet, despite progress identifying neural correlates of coordinate frame transformations in the auditory system of animal models, there is little systematic evidence demonstrating that non-human listeners experience sounds in multiple coordinate frames. This imposes a fundamental limitation on our ability to gain insight into the circuit mechanisms that support world centered hearing.

Egocentric and allocentric cognition has primarily been studied in navigation, where animals can use both strategies to navigate through environments (Burgess, 2006; Paul et al., 2009). However, most studies of navigation have focused on senses other than hearing (Fischler-Ruiz et al., 2021; Leutgeb et al., 2005; Muller and Kubie, 1987) and so leave open questions about the way in which non-human listeners perceive sound location. In contrast, studies of sound localization have adopted tasks that reveal key insights into neural processing, but do not specify the coordinate frame(s) in which sound space is defined (Bajo et al., 2019; Lomber and Malhotra, 2008; Wood et al., 2017). Here, we designed two tasks in which ferrets were trained to discriminate sound location in either world or head-centered space across changes in head direction to test the hypothesis that animals can localize sounds in multiple coordinate systems.

## Methods

### Animals

Subjects were seven pigmented female ferrets (0.5 to 3 years old, weighing 600-1100g) maintained in groups of two or more ferrets in enriched housing conditions, with regular otoscopic examinations made to ensure the cleanliness and health of ferrets’ ears.

Ferrets were trained either to report sound location in return for water rewards that were part of a daily minimum (60 ml/kg body weight) as part of a water regulation schedule in which animals also received supplementary wet mash made from water and ground high-protein pellets. Animals were water-regulated on a maximum of 50% of days, and were trained / tested in series of morning and afternoon sessions on consecutive weekdays, with rest weeks without testing at frequent intervals. The weight and water consumption of all animals was measured throughout the experiment.

All experimental procedures were approved by local ethical review committees (Animal Welfare and Ethical Review Board) at University College London and The Royal Veterinary College, University of London and performed under license from the UK Home Office (Project License 70/7267) and in accordance with the Animals (Scientific Procedures) Act 1986.

### Experimental Apparatus

Testing was conducted in a double-walled sound proof acoustic chamber (IAC Acoustics: 1.6m wide, 1.6m long and 1.1 m height) lined with 45 mm sound-absorbing foam and dimly illuminated using white LED striplights. Within the chamber, ferrets performed psychoacoustic tasks inside a custom-built, circular behavioral arena (0.5m radius) covered by plastic mesh to prevent animals escaping. The arena was surrounded by a ring of twelve speakers (Visaton FRS8) positioned at 30° intervals, 55 cm from the arena center (**Fig. 1A**). Below each speaker were also positioned lick ports that could detect the presence of animals via infra-red reflectance sensors (Optek OPB710), and deliver water via solenoid control (Flo-Control, Valeader).

**Figure 1.**
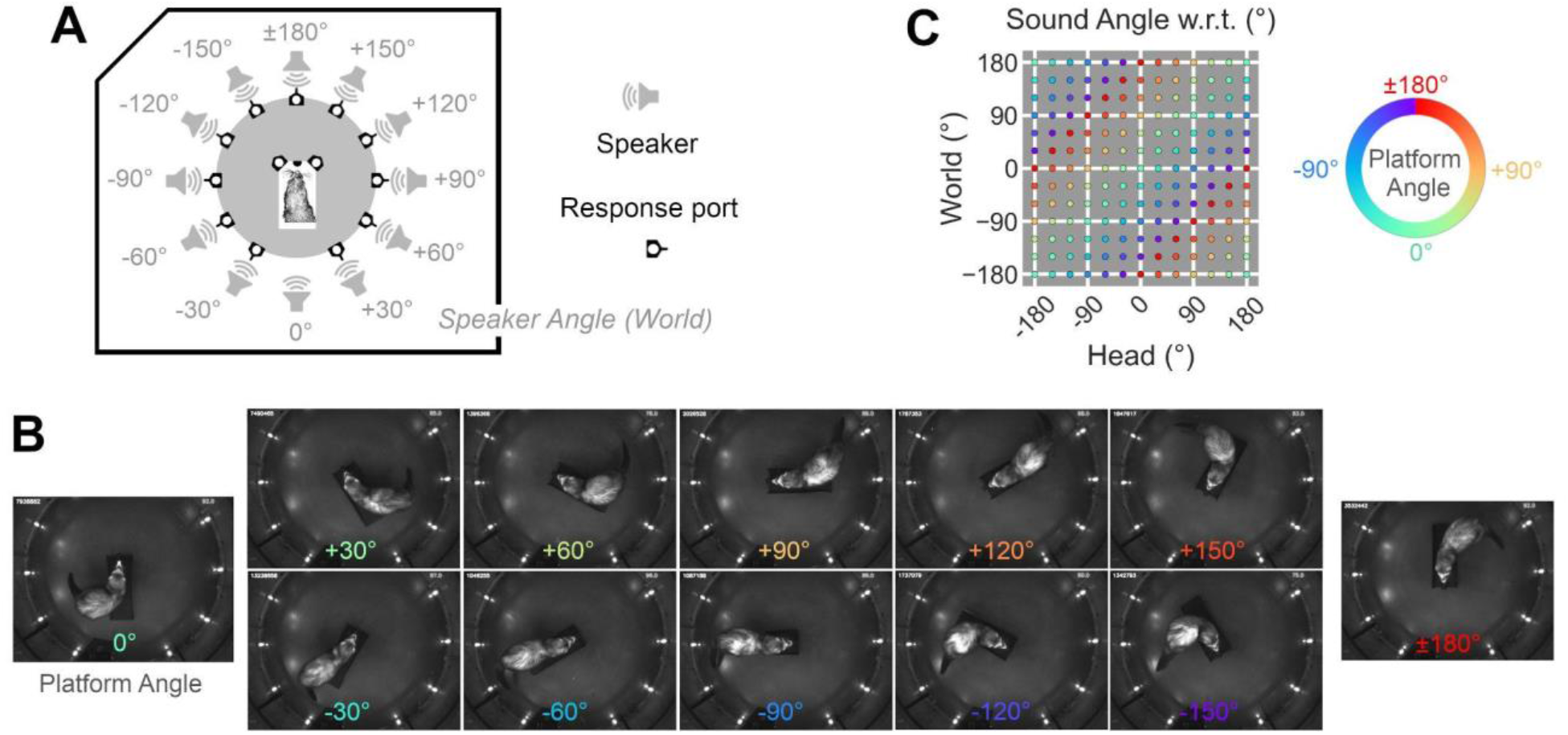
**A.** Task arena in which ferrets approached the center of a speaker ring to initiate presentation of a 250 ms broadband noise burst from one of twelve speakers. Values indicate speaker angle relative to the arena (world) coordinate system. **B.** Infra-red images showing one ferret at the center spout as platform angle is varied. Response ports around the arena periphery contain IR emitters and are thus highlighted here, but were not illuminated within the visible spectrum during testing. Values show platform angles (and thus head directions) within the world. **C.** Dissociation of speaker position in head and world-centered space that occurs with platform rotation.

At the center of the arena was a 3D printed platform from which animals initiated trials. The platform consisted of a 30-cm long section on which ferrets could align themselves, and an array of three lick ports (center, left and right) that detected nose-pokes and channeled water delivery. The platform was offset, such that the center of the ferret’s head was at the center of the test arena (and thus surrounding speaker ring) when visiting the center port on the platform. Between test sessions, the platform could be rotated at 30° intervals about the arena center, allowing us to orient animals throughout 360° within the test arena (**Fig. 1B**). By changing the platform angle and presenting sounds from different speakers within the ring, we could thus independently vary the position of sounds in the world and relative to the ferret (**Fig. 1C**). Here we use the term ‘platform angle’ to denote the orientation of the center platform relative to the arena, independent of any sound location in any coordinate system.

The sound-proof chamber in which experiments were conducted was dimly lit throughout (~10 to 20 lux), and visual landmarks within the test environment were limited to the 11 peripheral reward response ports (see below for use in each task) and the doors to the test arena. No specific visual landmarks indicated the east and west ports but the door at the front of the chamber, and the ‘missing’ rear response port at 0° provided visual anchors to the animals orientation within the environment. However, because ferrets’ vision is not fully panoramic, these landmarks were not visible at all platform angles, and thus provided unreliable visual cues. (The visual field of ferrets covers ~270° (Williams, 2012), while the range of angles within which each landmark fell covered 360°; with even the largest landmark [the arena door] subtending > 45° of the visual field). Although testing was performed in a double walled sound-proof chamber, we also presented a low level of background noise (45 dB SPL) from all 12 speakers continuously throughout each session to mask any variations in room noise.

### Task Design

Animals were trained in either a world-centered, or head-centered sound localization task. Both tasks had a two-choice design, in which animals were required to discriminate between a pair of sound sources from the speaker ring. On each trial, the animal would initiate presentation of a single 250 ms broadband noise burst from one speaker to which she could then respond. Broadband noise bursts were between 60 and 63 dB SPL, generated afresh on each trial.

To initiate sound presentation during each trial, ferrets were required to approach the central lick port on the platform and hold her head at the center of the arena. Prior to sound presentation, the subject was required to hold at the port for a variable delay (from 0.2 to 0.7 s), and trials would only be successfully initiated if the animal remained in position through the full duration of the sound. This hold period ensured that animals had a stationary head position during sound presentation (Dunn et al., 2021) and thus minimized any available dynamic localization cues. To encourage subjects to hold at the center port, rewards were delivered at this location on randomly allocated trials with a variable probability (upto 10% of trials). If an animal failed to wait for the full hold time, the trial was not initiated and the animal was required to release and reengage the center port to try again.

Following stimulus presentation, animals could respond by visiting one of two response ports based on the rules of the specific task (see below). Correct responses were rewarded with water; errors were signaled by presentation of a brief tone (500 Hz, 100 ms) and led to a short timeout (between 1 and 5 seconds) in which animals could not initiate further trials. Incorrect responses also led to correction trials, on which the same stimulus was presented. On a small proportion of trials (~10%), probe sounds were presented from untrained locations; responses to probe sounds were always rewarded. Animals were required to respond within a set time window (5 minutes) however animals rarely failed to respond, except when the ferret was saited and lost interest in performing the task, at which point the session was ended.

#### World-Centered Task

Ferrets (n=3: F1701, F1703 and F1811) were trained with sounds from either the North or South of a test chamber and required to report speaker location by visiting response ports with fixed locations in the chamber (East or West respectively, **Fig. 2A**). The subject was required to perform this North/South discrimination across rotations of the central platform (**Fig. 2B**). We also trained a fourth ferret with a different pair of world-centered locations (F1902: Respond West for sounds from −150° North-West vs. respond East for sounds from +30° South-East, **Fig. 2C**).

**Figure 2.**
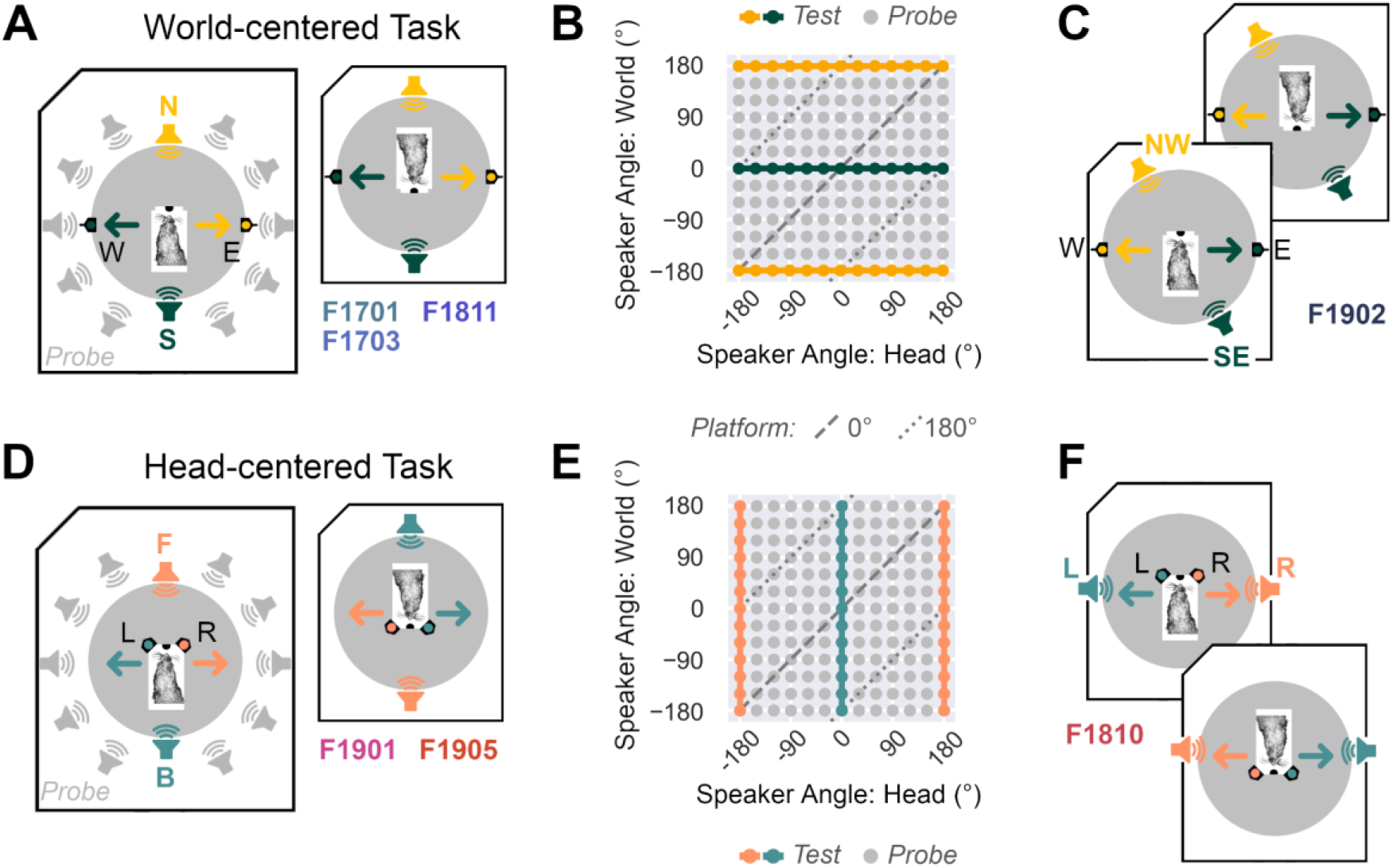
**A**. World-centered task in which subjects approached the center of a speaker ring to initiate presentation of a 250 ms broadband noise burst from a speaker either at the North (N) or South (S) of the arena, or later from probe speakers around the remainder of the arena (gray) by responding at East (E) or West (W) response ports. Arrows show the position of correct responses, which remained constant as the central platform was rotated. F-numbers (F1701 etc.) refer to ferrets trained in the North-South discrimination. **B.** Dissociation of speaker angle relative to the head and speaker angle in the world as the platform angle was rotated at 30° intervals. In addition to test sounds, probe sounds were also presented from untrained speaker locations on a random subset (10%) of trials. **C**. Variant of the task for an additional ferret (F1902) in which we altered the world-centered locations to be associated with each response. **D**. Head-centered task in which subjects discriminate 250 ms broadband noise bursts from a speaker either at the Front (F) or Back (B) of the head, by visiting response ports either to the left (L) or right (R) of the head. Arrows show the direction for correct responses. **E**. Dissociation of speaker angles in head and world-centered space as platform angle was rotated. **F**. Variant of the task in which we altered the head-centered locations associated with each response for one ferret.

#### Head-Centered Task

Ferrets (n=2: F1901 and F1903) were presented with sounds from either directly in front of (180°), or behind the platform (0°) and required to report speaker location by visiting response ports on the platform (60° Right or Left of the central port respectively, **Fig. 2D**). To test this front-back discrimination across changes in platform angle, we presented stimuli from different speakers in the ring as the platform rotated so that test sounds were always at the same angle relative to the head (**Fig. 2E**). Subjects were thus required to discriminate head-centered sound location, across changes in sound position in the world. We also trained a third ferret (F1810) to respond Left for sounds to the left (−90°) and Right for sounds on the right (+90°) of the head (**Fig. 2F**).

### Training

We initially trained animals using a widely used approach-to-target sound localization design (Dunn et al., 2021; Parsons et al., 1999; Wood et al., 2017). Animals were first acclimatized to the test arena and then trained to associate response ports with water. Training in each task took place with the platform facing the same direction (North: 0°), with the exception of F1810 (head-centered task trained facing West: +90°).

#### World-Centered Task

To train ferrets in the world-centered task, we then presented subjects with repeating colocalised audiovisual stimuli (the noise burst, accompanied by a white LED with an interstimulus interval of between 100 and 250 ms) from the East / West response port locations surrounding the edge of the arena. Ferrets typically learn to approach such stimuli in return for water rewards within one to two weeks; initially responses at either port were rewarded, however error trials were introduced once animals had established a reliable pattern of initiating stimuli at the center platform and visiting response ports. The timeout duration following error trials was initially set to zero, and increased gradually as animals performed more trials with training. Through training, we also gradually increased the time that animals were required to hold at the center port before stimulus presentation.

Once the animal was accurately performing the baseline audiovisual localization, we progressively shifted the speaker presenting sounds away from the (e.g. East or West) response port to the final trained location (North or South) over a series of weeks. Over the same period, we also (i) lowered the LED intensity at the response port until it was completely absent and no visual stimulus was presented and (ii) reduced the number of stimulus repetitions to one. By the end of this stage, the animal was responding accurately to a single auditory stimulus (250 ms broadband noise). We considered animals to be fully trained and ready for testing if they could maintain good performance (typically ≥ 70%) under these conditions, over multiple days. The complete training process typically required three to six months.

#### Head-Centered Task

To train ferrets to discriminate sounds based on head-centered location, we introduced repeating noise bursts when the subject held at the center initiation port for a sufficient time. Noise bursts were presented from either left or right of the head, and responses at either port on the platform were initially rewarded. Once animals had learnt to reliably initiate and respond to sound presentation, errors were introduced so that animals were required to respond at the Left response port for sounds on the left, and respond at the Right port for sounds on the right. Across multiple sessions, we then shifted the sound source associated with Left and Right response port counter-clockwise until animals were discriminating sounds from in front (respond Right) or behind (respond Left). We then reduced the number of repeating stimuli to a single 250 ms noise burst, while also increasing the hold time required to initiate sound presentation, and increasing the timeout that resulted from error trials. Typically it required between two and three months to train animals to achieve >70% discrimination of a single sound presentation.

#### Distal vs. Proximal Responses

Although the training methods were similar, training in the two tasks required different coordination of behavioral responses at different scales; the head-centered task took place in peripersonal space in which the animal could respond with only head movements, and there was no need for further locomotion. In contrast, the world–centered task required animals to run to and from distal response ports on every trial performed. As each response required the animal to run 1 m, this accumulated across trials and training sessions, so that a ferret performing the world-centered task would run approximately 1 km more per week within the arena than a ferret performing the head-centered task. This additional movement and associated exploration of the test arena during training for the world-centered task may support the formation of allocentric representations, such as stable place cells in the hippocampus (Frank et al., 2004; Kentros et al., 2004) and thus enhance the world-centered framework in which sounds could be mapped.

### Testing

Once animals had completed training, we began rotating the central platform at which trials were initiated. Platform rotations were initially made at small angles (e.g. 30°) to ensure that animals could continue to initiate trials successfully, and then broadened to larger angles until the full 180° rotation was achieved. Animals were then tested at all platform angles (30° intervals, 360° range) in a pseudorandom order that aimed to test all directions on similar numbers of sessions (with the exception of the initial training orientation). Here, the platform was manually rotated between behavioral test sessions when the subject was not in the test chamber, and thus on each session, a subject completed many trials (typically 50 to 100 in world-centered localization, or 50 to 300 in head-centered localization) with the same platform rotation.

#### Probe Testing

To determine how animals thought about sound space beyond the locations of the two test sound sources (e.g. North and South, Front or Back), we also presented probe sounds from the remaining ten speakers in the ring that surrounded the arena. To avoid familiarizing animals to the location of probe sounds, or eroding the subject’s discrimination of test sounds, we only presented probe sounds on a maximum of 10% of trials in any session. Occurrence of probe trials and speaker location on each probe trial was pseudorandom, but always excluded the first 10 ten trials of each session that were reserved for test sound locations. Responses at either of the two response ports used in the task were rewarded.

#### Speaker Swaps

To exclude the possibility that world-centered discrimination of sound location was driven by any unknown acoustic properties of sound sources (e.g. notches in spectral output that we could not detect) speaker swaps were performed. This involved physical disconnection of two speakers from the stimulus generation hardware, with each speaker being reconnected in the location and to the input connections of the other.

### Data Analysis

All correction trials were excluded from data analysis.

#### Ferret Behavior: Discrimination of test sounds

To report overall performance discriminating sounds from test locations as percent correct, we combined data across multiple test sessions. As we had unequal sample sizes, with an oversampling of trials with the platform in it’s initial training orientation, and because there was individual variability in the number of trials performed by each ferret, we randomly subsampled performance on 400 trials without replacement for each animal and each platform direction. (This number was selected based on the smallest sample size in the project dataset). To ensure reliable measurements, we repeated (bootstrapped) resampling 100 times and reported the mean performance across samples. Responses to probe sounds were not included in this analysis.

For each ferret and platform angle, we also assessed the probability of observing task performance by chance using the binomial test, where the probability of success under the null hypothesis was 0.5. Task performance for these tests was defined as the mean number of correct trials performed across bootstrap iterations, rounded to the nearest integer. The total number of trials was fixed (n=400) for all comparisons.

#### Ferret Behavior: Responses to test and probe sounds

We measured the probability of making a particular response (e.g. to go West or Left) as a function of sound angle in head and/or world-centered space, using responses to both test and probe sounds. As the number of combinations of possible sound locations and platform angles was large (n = 144) and the probability of a probe sound being presented was low (10%), we had potentially few trials for some data points. To ensure equal sample sizes, we again subsampled our data using bootstrap resampling, in this case with replacement (3 trials per combination of head and world-centered sound angles, totaling 432 trials in total across all combinations).

To compare how strongly behavior was modulated by sound angle in different coordinate systems, we measured the variance in response probability associated with sound angle in a specific coordinate system. Variance was calculated as the sum of squared differences between response probability at each sound angle (*x_θ_*) and mean response probability across sound angles (*x̅*), normalized by the number of sound angles (*n* – 1):

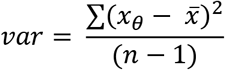

#### Behavioral Models

To understand the generative processes underlying animal behavior, we compared performance of ferrets with simulations from different model systems, and subsequently fitted parameters of these models to behavioral responses to identify the most likely explanations of our results.

##### World-centered task: Models of world-centered responses

To understand performance in the world-centered task, we began with two models that generated responses at a fixed location in the world (the West response port) as a function of sound angle in head or world-centered space (*θ*) using sinusoidal function with three parameters:

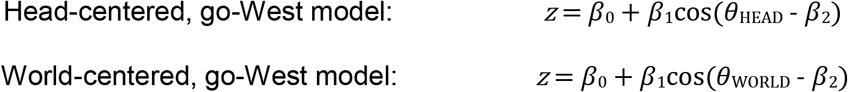

Here, the response variable (*z*) reflects an activation value associated with making a response at the West response port. Activation values for the alternative response (East) were defined as: *1 – z*. The model parameters were chosen to mirror the subject’s response bias (vertical shift: *β*_0_), sensitivity to sound angle (amplitude: *β*_1_) and the sound location at which a response was most likely (horizontal shift: *β*_2_). To convert activation values into response probabilities from which individual trial responses could be drawn, we used the softmax function with a variable inverse temperature parameter (*β*_tav.temp_) that reflected the extent to which the decision making was deterministic (*β*_inv.temp_ → ∞) or random (*β*_inv.temp_ → 0).

##### World-centered task: Models of head-centered responses

We also considered several models that generated responses in a head-centered coordinate system using some fixed offset relative to the head-centered sound location. For example, in simulations, we considered a model that responded at a 90° location relative to a presented sound (i.e. that went to the listener’s right when a sound was in front, or left when a sound was behind). Here, the offset between response and sound locations represents a parameter (*β*_OFFSET_) in the model:

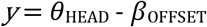

The response variable (y) reflects the location of the output response in head-centered coordinates, which can then be converted to world-centered coordinates by subtracting the angle of the head in the world (determined by the platform angle). However, without further modification, such models would predict responses on probe trials at world-centered locations for which no response port existed in the task (null locations). We therefore added three alternative strategies through which such models might compensate, allowing them to convert responses at null locations into responses at active East or West responses. As logical statements, these strategies took the form ‘if no port exists at target response location, then...’:

Strategy 1: Guess randomly
Strategy 2: Go to the nearest active port, or guess if equidistant
Strategy 3: Distribute responses over available ports, weighted by the relative distance between target response and active ports

These strategies allowed us to compress the model outputs into activation values for East or West response ports, which we then converted into probabilities using the softmax function with the same inverse temperature parameter (*β*_tav.temp_) described for earlier models.

##### Head-centered task: Models of head-centered responses

To model performance when discriminating sounds in the head-centered task, we simply adapted the same model format initially used for the world-centered task. However, rather than model East/West responses, we switched the response space to give the probability of responding at Left or Right:

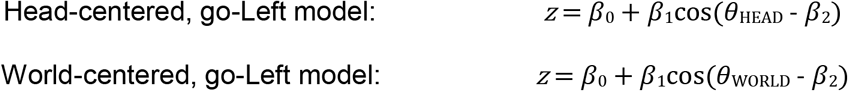

Here, the response variable (*z*) reflects an activation value associated with making a response at the Left response port. Activation values for the alternative response (Right) were defined as: *1 – z*. The parameters in the model have similar biological interpretations as presented above for the world-centered task, and we used the same softmax function with inverse temperature as an additional parameter to capture the determinism of decision making.

#### Simulations

To obtain performance metrics (percent correct) for models with known parameters, we simulated presentation of stimuli from all test locations that were used in experiments with ferrets. Response probabilities were used to define a multinomial distribution (NumPy)(Harris et al., 2020) from which we drew 10^3^ values, each of which represented single trial responses in simulations. We then summed the number of responses that would be correct under the rules of the relevant world-centered or head-centered task.

For simulations of the world-centered task, using models generating responses in a world-centered system, we used the following parameter values: *β*_0_ = 0.5, *β*_1_ = 0.3, *β*_2_ = 0 and *β*_inv.temp_ = 2. For simulations of the world-centered task using models generating responses in a head-centered system, we set parameters as: *β*_OFFSET_ = −90, and temp = 1. For simulations of the head-centered task, using models generating responses in a head-centered system, we set parameters as: *β*_0_ = 0.5, *β*_1_ = 0.3, *β*_2_ = 0 and temp = 2.

#### Models Fitting

To find model parameters that most closely approximated ferret behavior, we used 20-fold cross validation to split data into train and test sets on which we performed model fitting and evaluation respectively. Prior to fitting, we combined data from animals with the same training conditions (world-centered task: F1701, F1703 and F1811, head-centered task: F1901 and F1905). We then flattened the distribution of sound angles in head and world coordinates in the training dataset by randomly selecting a fixed number of trials (n=10) for each unique speaker location in head and world-centered space (n=144).

Across varying parameters of each model, we minimized the negative log likelihood between training data and model output using the *fmincon* solver (Matlab, R2018b, MathWorks Inc.)(Wilson and Collins, 2019). Initial values were selected randomly within the bounds defined for each parameter (**Table 1**). Initialization and minimization was repeated 20 times for each fold to avoid local minima and identify the parameters that resulted in the lowest negative log likelihood.

**Table 1:**
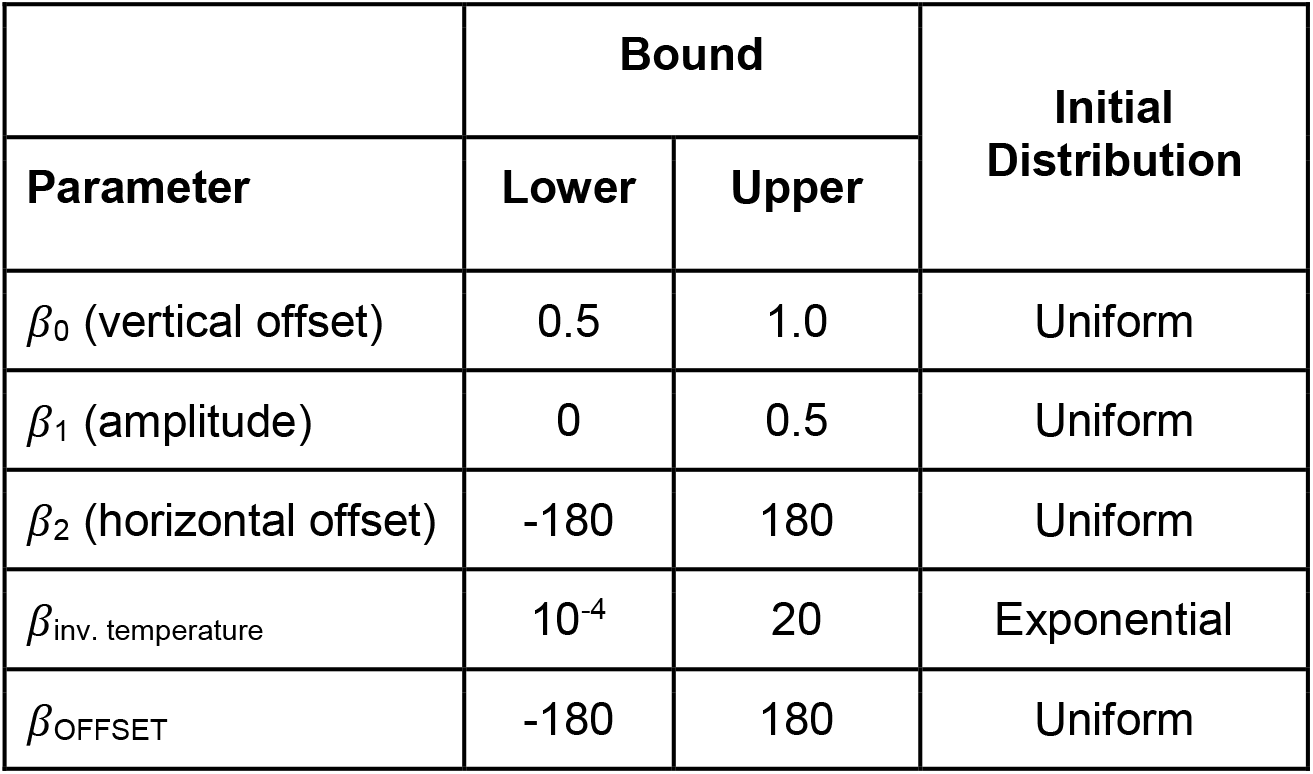
Parameter boundaries for model fitting.

For each model, we evaluated the parameters that gave the lowest negative log likelihood values on each fold. Evaluation used the sound angle in the modeled coordinate system, together with these parameters, to predict behavioral responses (i.e. whether the subject would go East/West or Left/Right) on single trials in held-out test data. Evaluation performance was measured as the percentage of trials on which we correctly predicted responses made by ferrets, and was used to compare the validity of different models.

#### Ferret Behavior: Head tracking

Some of the models we considered in our analysis of the world-centered task predicted behavioral responses at (null) locations other than East or West port, for which animals would then have to learn to compensate. To test whether ferrets actually attempted to respond at null locations, we measured the paths that animals took while responding to probe sounds by tracking head movements on each trial using DeepLabCut (v.2.2.3) (Mathis et al., 2018).

We selected trials from the first 10 sessions that ferrets (F1701: n = 31 probe trials, F1703: n = 49 and F1811: n = 39) were tested with probe trials to minimize any potential effects of learning. For each session, we labeled the head position in 20 images, as well as position of the nose, shoulders, spine and tail for each ferret and session (giving a minimum training set of 200 images per ferret; in the case of F1811, we added an additional 200 images when refining a trained network). We then trained a ResNet-50 based neural network for a minimum of 200,000 iterations starting with the default parameters. As the video system used to monitor animals was upgraded through the project, we trained separate networks to track F1811, and combined data from F1701 and F1703. We then used the network to analyze all frames within the dataset. Validation errors for test and training data of 5.37 pixels and 2.05 pixels respectively (image size was 460 by 640) for the network trained with videos from F1701 and F1703, and of 3.27 pixels and 2.21 pixels respectively (image size was 360 x 640) for the network trained on videos of F1811.

To analyze the paths taken by animals when responding to sounds, we cross-referenced the frame times captured in each video with behavioral timestamps captured by the data acquisition system, and selected frames between trial initiation (defined as the end of sound presentation) and trial completion (defined as the point at which the animal responded at one of the valid response ports). We used a p-cutoff of 0.1 to exclude any frames for which X or Y coordinates were missing for data visualization, and a p-cutoff of 0.5 to exclude any trials with missing data when comparing path lengths. Path lengths were taken as the sum of changes in position between trial initiation and response time. We then compared the path lengths taken on test and probe trials using a general linear mixed model (GLMM) with ferret as a random effect.

In a supplementary analysis to validate the alignment of platform angle and head direction in the head-centered task, we also labeled landmarks (left ear, right ear, nose and midpoint of the head) on a small sample of frames from randomly selected trials (n = 87). This sample was sufficient to detect minor offsets in head direction and so we did not expand the analysis to consider further data, nor train DeepLabCut models to track landmarks.

#### Ferret Behavior: Reaction time analysis

When considering the possibility that ferrets might respond at null locations in the world-centered task, we reasoned that such irrelevant responses would cost animals time in completing trials. In particular, we might expect animals to be slowest when least sure about the appropriate response to make; for example, when the intended response is equidistant with the two available ports. If true, reaction times should increase as animals distribute their responses more evenly between East and West response ports (i.e. the probability of responding West approaches 0.5).

To analyze reaction times, We first compared reaction times with unmodified response probability (p) for visualization purposes; we then adjusted response probabilities to give the distance from chance performance (*p*’) as:

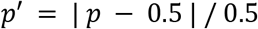

We then measured the association between reaction times and adjusted response probabilities, by fitting a GLMM with reaction time as the response variable, adjusted response probability as fixed effect and ferret as a random effect. GLMMs were performed in python using the statsmodels library (v0.13.1, www.statsmodels.org).

## Results

We tested if ferrets could discriminate between two sound locations that were either (i) fixed in the world, while varying relative to the head (**Fig. 2A-C**, n = 4 ferrets: F1701, F1703, F1811 and F1902) or, (ii) fixed relative to the head while varying in the world (**Fig. 2D-F**, n = 3 ferrets: F1810, F1901 and F1905).

### Discrimination of sound sources fixed in the world

Ferrets (n=4) successfully learned to discriminate between sound sources with fixed locations in the world. **Figure 3A** shows the behavior of each ferret trained in the task, with performance being better than chance at all platform angles (n = 12 angles, 400 trials per angle: range = 66.6 to 85.9% vs. 50% chance; mean performance of each ferret: F1701 = 74.6%, F1703 = 76.3%, F1811 = 80.3%, F1902 = 77.3%). Binomial tests confirmed that the probability of performance arising by chance was significantly low at all platform angles for each subject (Bonferroni corrected, p < 0.001). Similar results were obtained when considering the first ten trials in each session after platform rotation (**Fig. 3B**), indicating that accurate performance did not simply result from rapid relearning of the task after every change in platform angle.

**Figure 3.**
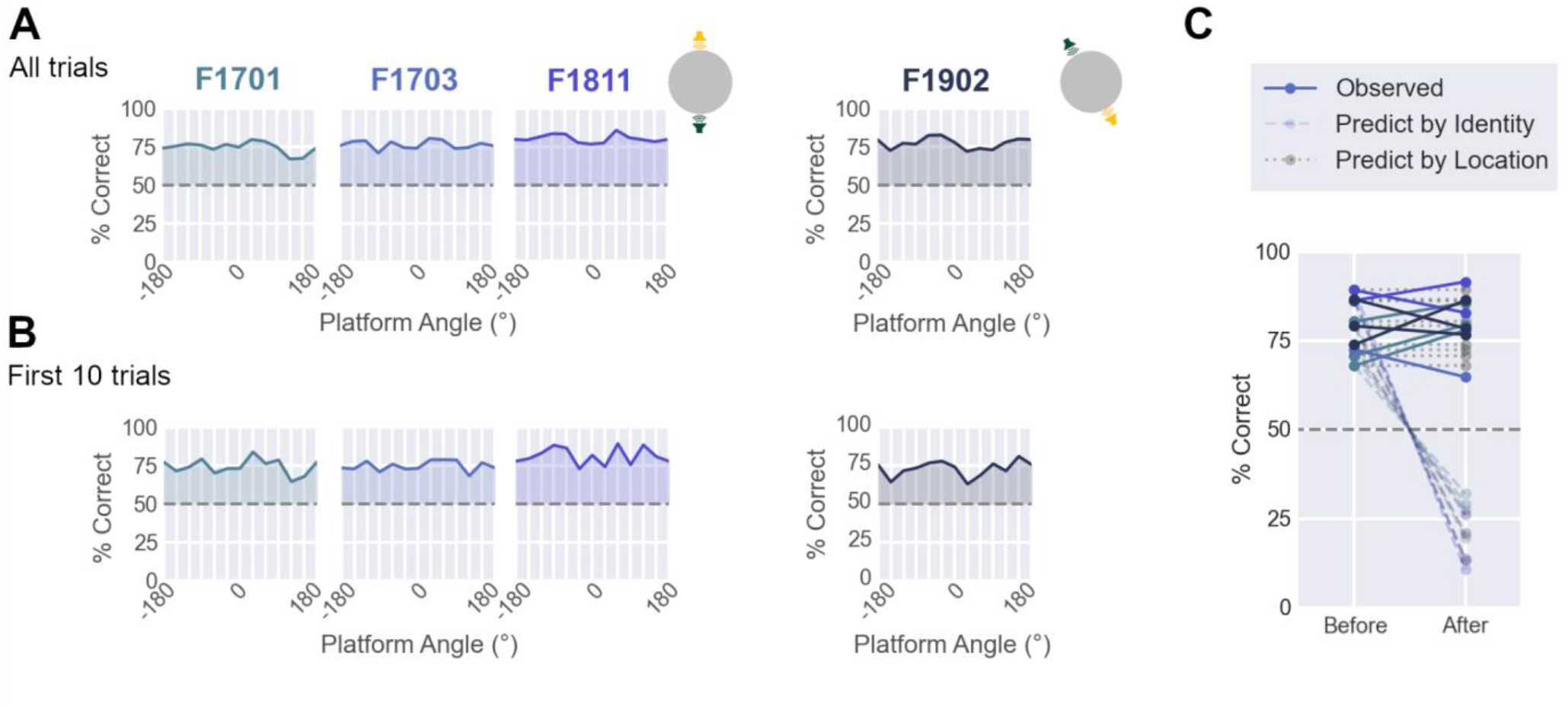
World-centered task performance. **A**. Performance discriminating sounds at trained locations for each ferret as a function of platform angle (n=400 trials per platform angle). Data shown as mean percent correct across bootstrap resampling (n=100 iterations). Dashed lines show chance performance (50%). Insets show the training configurations (either North vs. South [F1701, F1703, F1811], or SouthEast vs. North-West [F1902]). **B**. Performance measured only from the first ten trials of sessions after platform rotation (n = 50 trials per platform angle). Data shown as in A. **C**. Performance on sessions immediately before and after swapping the speakers at test locations. Observed data (full lines) is compared to predictions made if animals were responding based on speaker identity (dashed lines) or sound location (dotted grey lines). Predictions based on speaker identity were made by subtracting performance before swap from 100%. Predictions based on sound location were simply the same performance before and after swap.

To ensure that animals were using speaker location rather than speaker identity to solve the task, we also swapped the specific sound sources used at test locations (e.g. North and South, or North-West and South-East). Swapping speakers did not affect ferrets’ ability to discriminate world-centered sound location, as performance remained consistent before and after swaps (mean ± s.d. change = 1.906% ± 7.670%, **Fig. 3C**). Thus ferret behavior was driven by the location of the sound in the arena, rather than the specific speaker from which sounds were presented.

### Generalization to probe sounds

Our results show that ferrets can discriminate between the location of two sound sources that are fixed in the world. We next asked whether such discrimination reflected learning of specific associations, or if ferrets responded as a continuous function of sound location in a specific coordinate system. To address this, we measured ferrets’ responses to probe sounds presented on a random subset (10%) of trials from speakers at untrained angles (n=10) around the test arena (**Fig. 2B** and **2E**).

For animals trained with sounds that were fixed in the world, probe testing revealed that the probability of each ferret making a specific response (e.g. to visit the West response port) was a continuous function of sound angle in the world (**Fig. 4A**). In contrast, changes in sound angle relative to the head had no consistent effect on behavior (**Fig. 4B**). This graded pattern of responses was observed regardless of whether animals were trained to visit the West port to indicate sounds from the South (F1701, F1703 and F1811) or from the North West (F1902). The same coordinate frame-specific modulation was apparent in the joint distribution of responses, where behavior was most strongly influenced by the position of sounds in the world (**Fig. 4C**).

**Figure 4.**
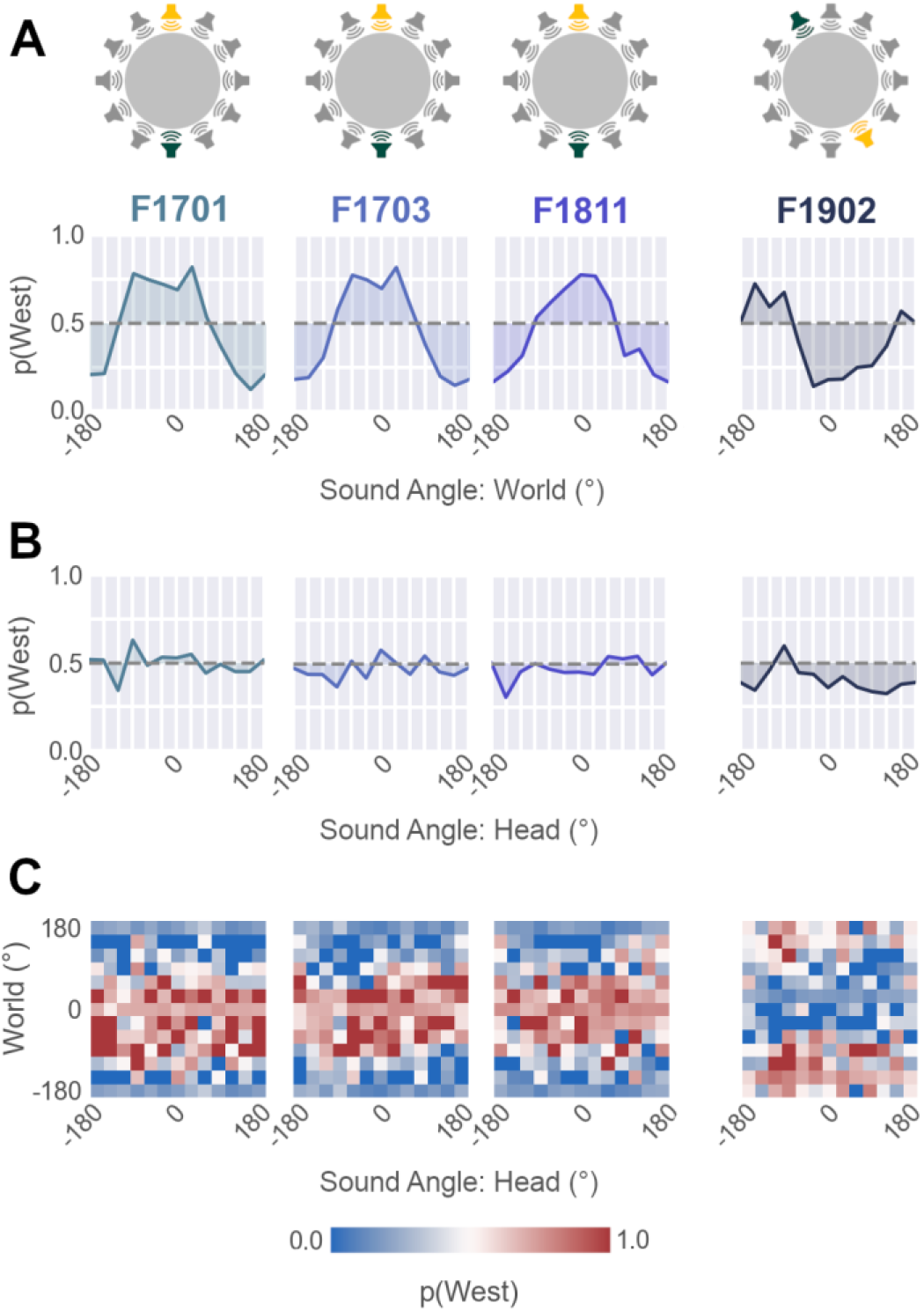
**A-B** Probability of responding at the West response port for test and probe sounds as a function of sound angle in the world (**A**), or relative to the head (**B**) (n=36 trials per angle) in ferrets trained in the world-centered task. **C.** Response probability as a joint function of head and world-centered sound angle (n =432 trials over 144 locations).

We quantified modulation of behavior by sound location in each coordinate frame by calculating the variance in response probability across sound angles in head or world-centered space. Higher variance indicates greater modulation of behavior by sound angle in the respective coordinate frame. In animals trained with sounds fixed in the world, we found larger variance associated with sound angle in world-centered than head-centered space (Variance: World vs. Head; F1701: 0.067 vs. 0.005, F1703: 0.066 vs. 0.004, F1811: 0.052 vs. 0.004, F1902: 0.043 vs. 0.007).

### Simulating world-centered task performance

Our data from the world-centered task show that ferrets can discriminate between two sound locations that are fixed in the world, across the changes in sound angle relative to head that occur when the platform is rotated. Moreover, when presented with probe sounds from untrained locations, animals respond as a continuous function of sound angle in the world. These results are consistent with the suggestion that ferrets can localize sounds in a world-centered coordinate system; however, to confirm this interpretation we must first consider alternative strategies that might produce similar behavior (**Fig. 5**).

**Figure 5.**
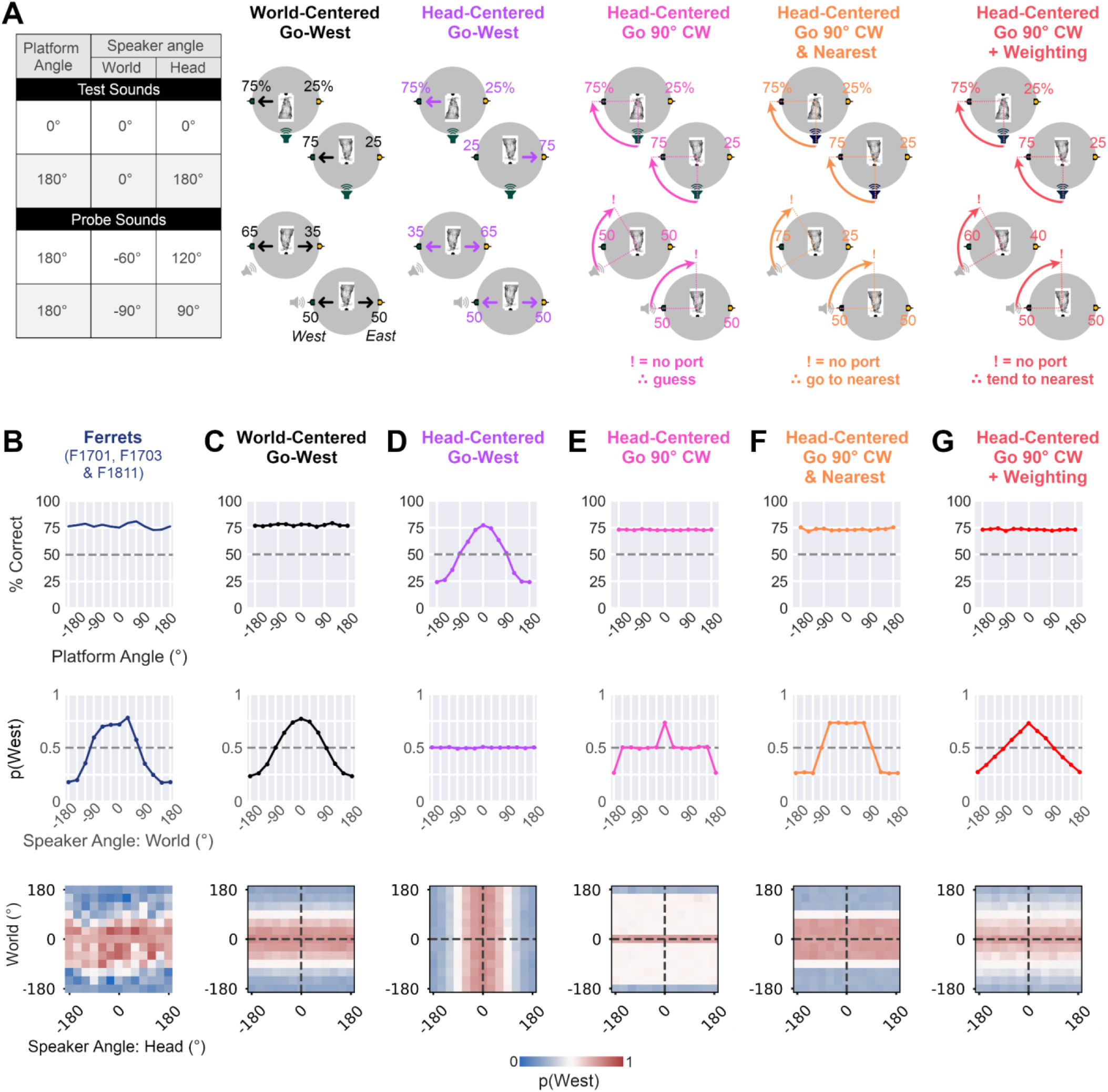
Models of world-centered task performance. **A**. Probability of responding at East or West response port under four example conditions in which platform angle and speaker location is varied. Values show probability of responding at East and West ports, expressed as percentage. Exclamation marks indicate trials for which the model would attempt to respond at an inactive (null) port. **B**. Performance of three ferrets trained with the same pair of world-centered sound locations (F1701, F1703 and F1811) in terms of overall accuracy (top row: % correct), probability of responding at the West port as a function of sound angle in the world (middle) and West response probability as a function of sound angle in head and world-centered space (bottom). **C-G**. Corresponding predictions from simulations of each model (see methods for details of model parameters in each simulation).

We began by contrasting behavior of ferrets trained with the same world-centered locations (**Fig. 5B**: F1701, F1703 and F1811) with simulated performance of two models that linked responses at the East or West response ports to sound location in a specific coordinate frame. A world-centered model generated responses as a continuous function of sound angle in the arena (**Fig. 5C**). Across platform rotations, this model successfully discriminated between test sounds at North and South locations in the arena and produced a pattern of responses to probe sounds that was similar to ferrets. In contrast, a head-centered (specified here as the ‘head-centered, go-west’ model, **Fig. 5D**) only performed well when the platform was oriented at specific angles, and did not replicate the behavior of ferrets across platform angles.

We then considered alternatives of the head-centered model that responded not in world-centered space, but rather in head-centered space. That is to say, rather than determining whether the listener responded at the East or West response port, these models determined whether the animal made a specific head-centered response; here, to respond at the port 90° clockwise of the head-centered sound location at which stimuli were presented. With this strategy, the model could successfully discriminate between test sounds at North and South speakers (**Fig. 5E-G**). However, such models alone could not explain responses to probe sounds, as a 90° clockwise response would take the animal to response ports that were not part of the task (null ports). For example, if a probe sound was presented at the South-West, the animal should respond at the North-West port, but this response port was not active in the task and the outcome of the trial was thus undetermined.

If a model predicted responses at null ports, we added one of three possible assumptions about the strategy the subject might take. In the first instance, we simply assumed that any target response other than East or West would lead to random guessing (**Fig. 5E**), but this failed to account for the structured response patterns shown by ferrets to probe sounds. We therefore extended the model to respond at the nearest available port, or guess if the target port was equidistant to East and West ports (**Fig. 5F**). This simulation (“Go 90° CW & Nearest”) generated a stratified response profile in which the model always responded with maximal probability at a particular port or otherwise at chance levels, and thus failed to mirror the graded response profile shown by ferrets. Finally we considered a strategy in which response probabilities for East and West ports were weighted by the distance between the target null port and the two active ports (“Go 90° CW + Weighting”). Although this model produced a graded response to probe sounds that was more consistent with ferret behavior than the other head-centered models (**Fig. 5G**), the equal spacing between response ports resulted in a linear relationship between world-centered location of probe sounds and response probability that contrasted with the sinusoidal profile shown by ferrets, and predicted by other world-centered models (**Fig. 5C**).

### Modeling behavior

Our simulations illustrate prospective patterns of behavior generated by different models with known parameters; however it is also possible to fit the parameters of each model to the observed behavior of animals and determine which system best captures task performance. To fit models to observed behavior, we used 20-fold cross-validation on data from ferrets trained to discriminate the same pair of world-centered sound locations (i.e. North vs. South: F1701, F1703 and F1811). Model parameters were fit to a training subset of the data, so that we could then evaluate how well each fitted model predicted single trial responses from the held-out test dataset.

The world-centered model that linked sound location in the world to responses at East and West ports best predicted the behavior of all ferrets (**Fig. 6**). Validation performance across folds (median = 65.6% correct) was higher than alternative head-centered models that responded in head-centered space by going to the nearest port (63.2%) or by weighting responses according to distance with active ports (63.8%). Other head-centered systems predicted animal behavior at near chance levels, including models that linked sound location directly to East and West responses (50.1%), and models that responded in head-centered space but guessed on probe trials (49.8%).

**Figure 6.**
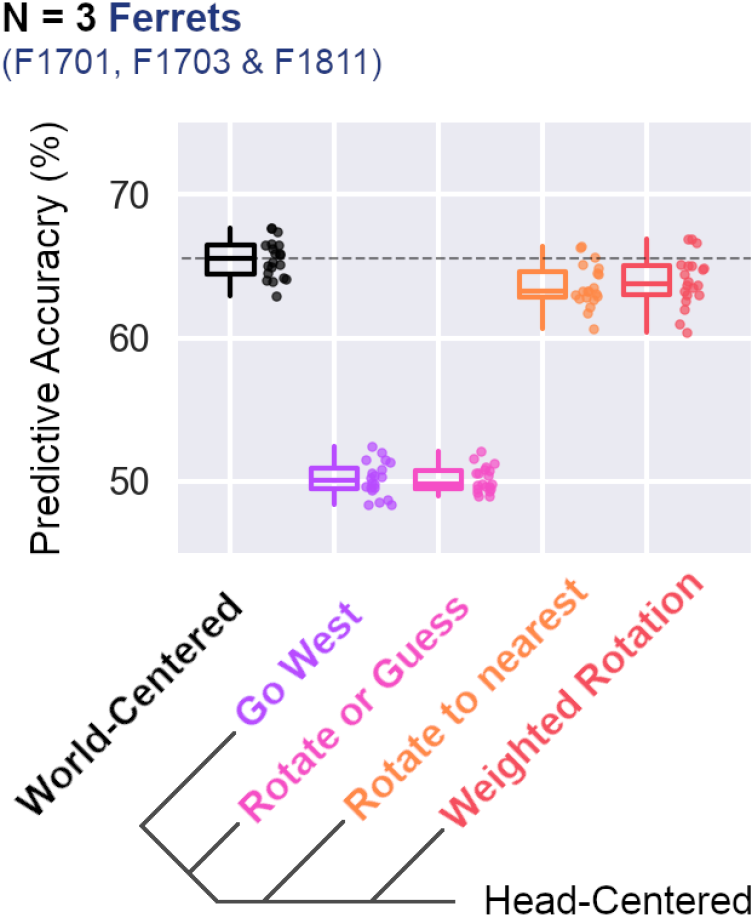
Model fit to animal behavior. Model validation performance showing accuracy in predicting single trial behavior from held-out data (20-fold cross-validation) collected from all ferrets trained to discriminate sounds from North and South locations (F1701, F1703, F1811). Box plots show median and interquartile range (IQR); individual data points show validation performance for each fold.

### Resolving competing models

The pattern of responses observed in ferrets performing the world-centered task was most consistent with a world-centered model in which animals used the position of sounds in the environment to respond at East or West response ports (i.e. world-centered sound localization). The next best model was provided by a head-centered strategy in which listeners responded at the nearest port that was a fixed rotation away from the head-centered sound location, with responses on probe trials resulting from a weighted guessing process. We note that our world-centered models used more parameters to achieve better performance predicting animal behavior (four compared to two). In contrast, head-centered models relied on additional assumptions about the strategy ferrets might use to respond on probe trials. Here, we tested these assumptions by looking at additional properties of behavior.

A key assumption of competing head-centered models was that animals would attempt to respond at inactive response ports, before using some strategy to redirect to the East or West ports that were active in the task. We therefore asked if ferrets ever tried to respond at ports other than the East or West by tracking the path of the ferret’s head on test and probe trials. Here, we focused on trials from the first 10 sessions in which probe sounds were presented (first ~40 probe trials for each ferret), so that the timeframe for learning any compensatory strategy was small and thus the chances of catching responses to inactive ports were maximized.

We found no evidence of responses to inactive ports by animals tested in the world-centered task. Tracking the head position of animals (F1701, F1703 and F1811; i.e. the ferrets presented in **Fig. 6**) did not show any notable deviation made by ferrets on probe trials, when compared to test trials, nor attempts by ferrets to respond at ports other than active East and West locations (**Fig. 7A-B**). Instead, response trajectories on probe trials appeared to be closely matched to those seen on test trials for each ferret. To quantify any differences in response trajectories, we compared the path lengths taken on probe and test trials using a general linear mixed model with ferret as a random effect; however, there was no effect of trial type (**Fig. 7C**, β = −8.64, p = 0.203). The trajectories of head movements were thus inconsistent with the suggestion that animals responded in head-centered space as suggested by alternative models of task performance.

A second assumption of alternative head-centered models was that responses on probe trials would be modulated by the distance between available East/West response ports and unavailable ports targeted based on head-centered sound angle. If this modulation reflected uncertainty about the correct response to make, we might expect reaction times to increase as animals became less certain (i.e. their probability of making a West response tended to 0.5). We did not see such a pattern in our data (**Fig. 7D**), and no significant relationship was observed when we assessed the association between adjusted response probability (where probabilities were adapted to give the distance from chance performance [p = 0.5] – see methods) and median reaction times using a general linear mixed model, with ferret as a fixed effect (**Fig. 7E**; β = −0.101, p = 0.265).

**Figure 7.**
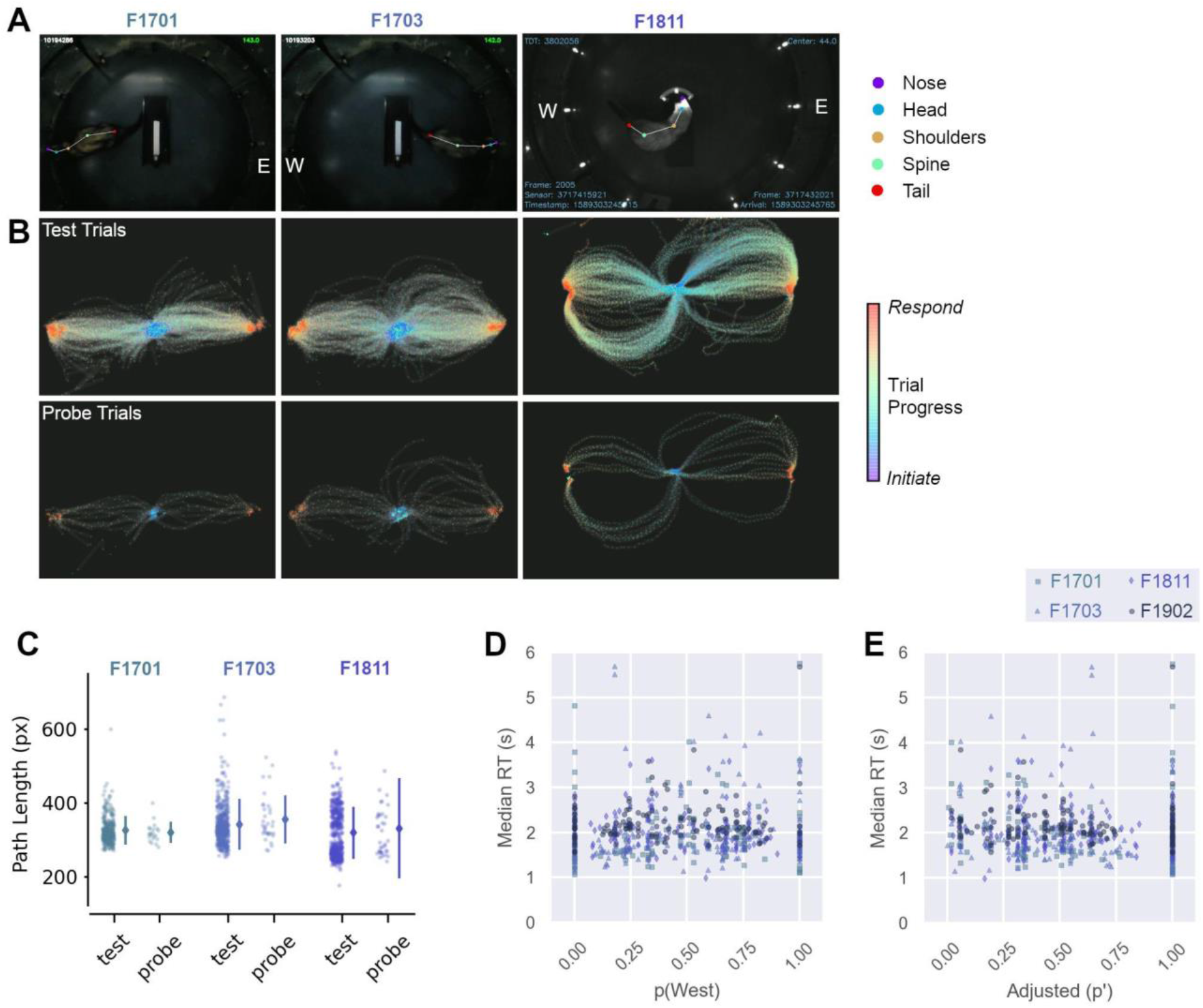
Head tracking and response time analysis. **A**. Screenshots showing tracking of head and body position using DeepLabCut. Response locations are labeled, for example East (“E”) and West (“W”) ports. **B**. Trajectories of head position during trials as animals responded to test and probe sounds. Data shown from responses in the first 10 sessions in which probe sounds were presented. Markers show positions on each frame; lines show linear interpolation between frames. **C.** Path lengths for data shown in B. Scatter plots show path lengths for individual trials, with lines showing mean and standard error for each ferret. **D.** Comparison of median reaction times (RT) with probability of responding at the West response port. Chance performance = 0.5. Median reaction times were calculated across trials for a given combination of speaker location in head and world-centered coordinates (n = 144 conditions per ferret). **E.** Reaction times as a function of adjusted response probability (p’)(see methods). Data shown as in E.

Together, reaction time and path analysis indicate that the assumptions of head-centered models were not borne out in the behavior of animals, and that ferrets did not have to adapt their behavior on probe trials as would be expected by such models. Thus, head-centered models provide a poorer account of task performance than those based on world-centered sound localization that more accurately predicted responses on both test and probe trials.

### Discrimination of sound source position relative to the head

We also trained a second group of ferrets (n=3) to discriminate sound locations that were fixed relative to the head (**Fig. 2D-F**). Subjects performed this task accurately when discriminating Front from Back (F1901 and F1905) or Left from Right (F1810) across platform rotations (**Fig. 8A**). Each animal performed above chance at all platform angles (range = 69.7 to 86.5% vs. 50% chance, mean performance of each ferret: F1810 = 73.4%, F1901 = 74.6%, F1905 = 82.7%). Binomial tests comparing observed performance to chance confirmed significant differences at all platform angles in each subject (Bonferroni corrected, p < 0.001).

**Figure 8.**
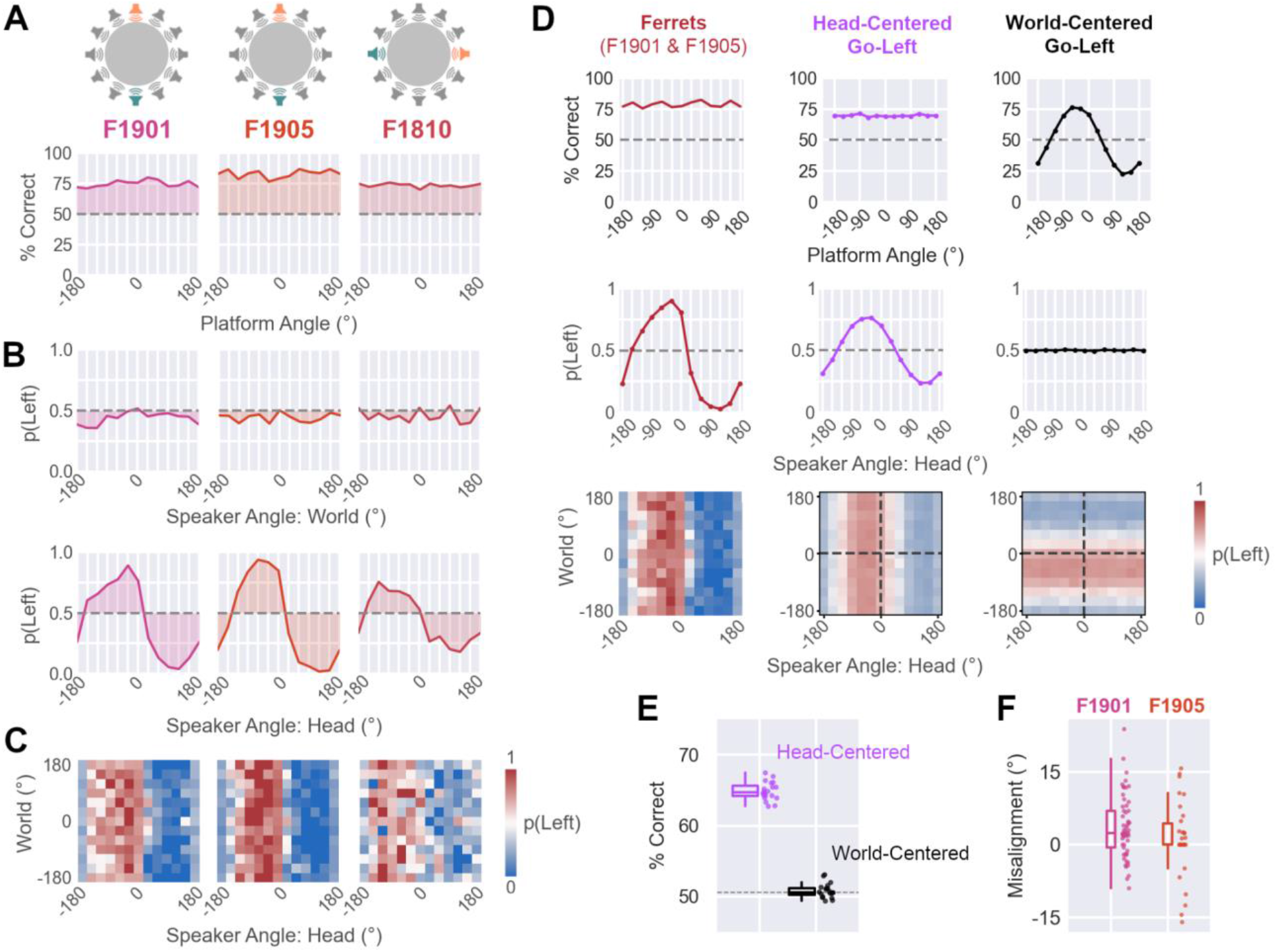
Head-centered sound localization. **A**. Performance discriminating sounds at trained locations for each ferret as a function of platform angle (n = 400 trials per angle). Data shown as mean percent correct across bootstrap resampling (n=100 iterations). Dashed lines show chance performance (50%). Cartoons show the training configurations (either Front vs. Back [F1901 etc.], or Left vs. Right [F1811]). **B**. Probability of making Left responses to test and probe sounds as a function of sound angle in the world, or relative to the head (n=36 trials per angle). **C**. Response probability as a joint function of head and world-centered sound angle, shown for individual animals (**C**, n=432 trials over 144 locations). **D**. Comparison of ferret behavior with simulated behavior of models that linked head or world-centered sound location to the probability of responding at the Left response port. Data shown for ferrets (F1901 and F1905) trained in Front-Back discrimination). Model parameters for simulations are given in Table 1 (see Methods). **E**. Model validation performance predicting single trial behavior from held-out data (20-fold cross-validation). Box plots show median and interquartile range (IQR); individual data points show validation performance for each fold. **F**. Distribution of offset values across trials (n=87) for animals trained to discriminate front and back sounds in head-centered space (F1901 [n=57] and F1905 [n=30]). In both cases, median offsets were significantly greater than zero (Wilcoxon rank sum, p < 0.001). Data shown as box plots indicating median and interquartile range, with individual trials shown as separate markers.

When tested with probe sounds at untrained locations, ferrets’ responses were strongly modulated by sound angle relative to the head, while sound location in the world had no comparable effect on response probability (**Fig 8B-C**). Larger variance was associated with sound angle in head-centered than world-centered space for all animals (Variance: World vs. Head; F1810: 0.003 vs. 0.043, F1901: 0.002 vs. 0.102, F1905: 0.001 vs. 0.140). Thus ferrets in this task reported the location of sounds as a continuous function of sound angle relative to the head, regardless of sound location in the world.

The behavior of ferrets in the head-centered task could be well captured by a model that linked head-centered sound location to the probability of making a response at the Left or Right port. Simulations with known parameters produce patterns of behavior that mirrored those observed in ferrets (**Fig. 8D**) and fitting model parameters to observed behavior allowed us to predict responses to sounds on a held-out dataset (**Fig. 8E**; median = 64.7% correct predicting responses for ferrets trained in Front-Back discrimination). In contrast, a competing model that linked world-centered sound location to responses at the Left port did not match with behavior in the head-centered task or predict held-out responses with substantially better performance than chance (median = 50.6%).

Finally, a notable feature of head-centered localization was the strong lateralization in response probabilities of animals trained to discriminate front vs. back sounds (F1901 and F1905). That is to say, the curves showing the association between head-centered sound angle and response probability in **Fig. 8B** are shifted leftwards, so that the animal is most likely to respond to sounds on the left by turning left (and vice versa on the right), even when trained to report sound in front at the left port. This behavior may have arisen because these animals slightly offset their head angle relative to the platform when initiating sounds: Video tracking revealed that this offset was small but statistically greater than zero (**Fig. 8F**: median = 2.22°, n = 87 trials, Wilcoxon rank sum test, p < 0.001) and aligned so that the rear speaker (to which animals responded by going Left) tended to fall more often on the left side of the midline. Likewise, the front speaker (to which animals responded by going Right) tended to fall more often on the right side. Across thousands of trials with test sounds, this small bias may have reinforced ferrets’ natural inclination to orient towards speakers.

## Discussion

Here, we established two behavioral tasks in which listeners were required to discriminate between the positions of sounds that were either fixed in the world (world-centered task), or fixed relative to the head (head-centered task). Subjects were then presented with probe sounds from untrained locations that measured whether listeners reported sound location as a continuous function of world or head-centered space. We found that ferrets could learn to perform either task, and that they responded as a function of sound angle in the task-relevant space.

To model task performance, we considered the response patterns of different head and world-centered systems; In the world-centered task, animal behavior was best captured by a model that linked sound location in the world to East and West responses. Alternative models based on responses in a head-centered system also mirrored animal behavior, but performed less well and required additional unmet assumptions to respond to probe sounds. In contrast, ferret’s responses in the head-centered task were best captured by a model that linked left and right responses to sound location relative to the head.

Together, our data thus suggest that ferrets can access information about the position of sounds in multiple coordinate systems, including sound location in the world, across variations in head-centered sound angle.

The distinction between egocentric and allocentric reference frames centered on the head/body and external environment has been the topic of extensive study in cognitive neuroscience, where representations in each coordinate system may be difficult to disambiguate. Here, we aimed to devise tasks that could only be solved using one coordinate system in order to clearly delineate the psychoacoustic abilities of non-human listeners. A key part of this design was the use of probe sounds, which in our analysis of the world-centered task, allowed us to contrast predictions from world-centered and head-centered models of sound localization. When designing experiments, we did not initially consider systems that would attempt to respond at inactive ports, and indeed there was no evidence that ferrets attempted to do so. However, the small difference in predictions of world-centered models and models that combined head-centered localization with a rotational heading rule suggests that improvements could be made to better isolate sound localization ability in a specific space.

In particular, our task design would be improved by requiring subjects to make a non-spatial response; for example, by using a temporal response dimension in which listeners are presented with sounds from multiple locations and required to respond only when a sound originates from a specific location. This approach would be similar to Go/No-Go designs that have been successful in other species (Amaro et al., 2021; Ferreiro et al., 2020). However, we wanted to avoid presentation of sound sequences in the current study, in case recent history of stimulus presentation affects neural processing of sound location (see below). Another alternative would be to use symbolic responses such as buttons with particular colors, shapes or symbols whose locations can be counterbalanced, and thus made irrelevant for task performance.

Our results indicate that non-human listeners can report sound location in head and world-centered space, but do not show whether the observed behavior is simply learnt, or reflects the ferret’s natural sound perception. Clearer insight on this issue could be gained by observing task performance when the opportunity for learning is absent. Such conditions arise on the first trial after rotating the platform by 180°, where the animal has yet to receive feedback about which coordinate system is task relevant. Our data are limited as we did not conduct these switch tests cleanly; instead we introduced small platform rotations at first to gauge their impact on animal behavior, which turned out to be minimal. However, this gradual introduction also gave animals the opportunity to learn about the task and thus prevented a clear test of the animals’ naive response. Future tests of world and head-centered sound localization should therefore build on the current results by conducting zero-shot tests of spatial generalization to gain a clearer picture into the coordinate frames animals inherently use without behavioral training.

Access to multiple sound spaces also raises the question of how flexible spatial representations are: Can ferrets switch rapidly between coordinate systems, as humans do when shifting between egocentric and allocentric descriptions of object location, or does switching require many trials for longer-term learning to take place? Cueing animals to locate sounds in specific spaces on each trial (Stoet and Snyder, 2003) offers one approach to address such questions in the future.

Our data show that ferrets could report the location of sounds in the world across changes in head pose, but the cues that subjects use to construct representations of the world itself remain to be determined. Key candidates for mapping include the visual and somatosensory landmarks involved in navigation, and integration of head direction into sensory processing in the hippocampus, entorhinal and retrosplenial cortex (Alexander and Nitz, 2015; Fyhn et al., 2007). It would therefore be valuable to test whether similar features are also critical for mapping sounds into the environment. Although we did not manipulate salient visual landmarks to study remapping, ferrets clearly knew the position of key features of the environment; most obviously the entrance to the arena, at which they would wait at the end of each session. By systematically varying environments (for example by having a moveable door or rotating arena), it may be possible to induce predictable shifts in world-centered sound localization that reveal the key anchors that ferrets use.

The conclusion that ferrets can report world-centered sound location despite changes in sound angle relative to the head tallies with observations in other species. Cats can update spatial judgements of sound location with proprioceptive and motor information during ongoing head and pinna movements (Ruhland et al., 2015) and gerbils can identify a sound source based on its position in the world (Amaro et al., 2021; Ferreiro et al., 2020). Our results extend these findings by fully dissociating sound localization in head and world-centered space to reveal access to continuous representations of sound location in multiple spaces. More broadly, and consistent with the egocentric coordinate frame transformations observed across the primate brain (Caruso et al., 2021; Werner-Reiss et al., 2003), our data contribute to a growing view that reconfiguration of sound space is a fundamental function of the auditory system.

How coordinate frame transformations take place within the auditory system is an open question. It is known that auditory cortex plays a key role in sound localization (Lomber and Malhotra, 2008; Wood et al., 2017), however the approach-to-target tasks used by earlier studies cannot shed light on whether auditory cortex is necessary for head, or world-centered localization, or both. In the case of world-centered localization, it will be critical to understand how non-auditory signals are integrated into neural networks involved in spatial hearing. Visual and vestibular systems offer information about head direction within the world that could support coordinate frame transformations, and interact with the auditory system at a variety of cortical and subcortical levels (Bizley et al., 2016; Wu and Shore, 2018).

Recordings from multiple brain regions during performance of tasks such as world-centered sound localization will be important in advancing our understanding how multisensory integration enables spatial hearing. In this regard, our task is optimally designed to streamline neural analysis, as: (i) Subjects must remain still on the central platform during sound presentation, which avoids complexities surrounding the effects of dynamic sound localization cues, or locomotor activity. And (ii) subjects are given only a single, short (250 ms) sound to discriminate, which means no potential interactions between sounds in sequences and minimal effects of stimulus history that might arise with continuous sound presentation. Characterizing the activity of neurons during sound presentation in the tasks developed here thus offers a way to clearly identify tuning to sound location in head and world-centered space.

The conversion between egocentric and allocentric representations is already the subject of intense scrutiny in navigation, where similar suggestions have been made for a network of interacting brain regions that includes retrosplenial and parietal cortex, and regions of the medial temporal lobe, including the hippocampus (Bicanski and Burgess, 2018; Wang et al., 2020). Despite the importance of these areas and the parallels in auditory and visual scene analysis, there is (to our knowledge) little known about the role of structures such as the hippocampus in spatial hearing. Recordings from echo-locating bats indicates that hippocampal function and auditory processing may be closely linked (Ulanovsky and Moss, 2007; Wohlgemuth et al., 2018), and in ferrets, hippocampal theta oscillations are widespread during approach-to-target sound localization (Dunn et al., 2021). Determining how the auditory system interacts with the medial temporal network, as well as parietal cortex, may thus provide important new insights into coordinate frame transformations in spatial hearing, and scene analysis more generally.

## Acknowledgements

We are grateful to Eleanor Jones, Zuzanna Slonina and Katarina Poole for help with data collection. An earlier report of results from a subset of the animals in this study was previously published in a peer-reviewed conference proceeding: Town & Bizley *Computational and Cognitive Neuroscience* Berlin 2019 doi: 10.32470/CCN.2019.1113-0.

This work was supported by the Biotechnology and Biological Sciences Research Council (BBSRC grant BB/R004420/1 awarded to SMT & JKB), Wellcome Trust (grant number WT098418MA awarded to JKB) and European Research Council (ERC Consolidator award [SOUNDSCENE] to JKB). This research was funded in whole, or in part, by the Wellcome Trust [Grant number WT098418MA]. For the purpose of open access, the author has applied a CC BY public copyright licence to any Author Accepted Manuscript version arising from this submission.

## Data Availability

All code and data associated with the project is available at: github.com/stephentown42/coordinate_specific_sound_localization

